# Sputum Proteomics reveals unique signatures linked to key outcomes in cystic fibrosis trials

**DOI:** 10.1101/2024.07.23.604741

**Authors:** Sian Pottenger, Dilip Nazareth, Dennis Wat, Bruno Bellina, Maike Langini, Martin Walshaw, Daniel R. Neill, Carsten Schwarz, Freddy Frost, Joanne L. Fothergill

**Author notes:** Correspondence: Jo Fothergill Department of Clinical Infection, Microbiology and Immunology, Institute of Infection, Veterinary and Ecological Sciences, University of Liverpool, Liverpool, United Kingdom and Freddy Frost Department of Cardiovascular and Metabolic Medicine, Institute of Life Course and Medical Sciences, University of Liverpool, Liverpool United Kingdom. These authors contributed equally (Freddy Frost, Joanne L. Fothergill). The authors have declared any conflicts of interest.

## Abstract

**Rationale:** Lung function (FEV1) and quality of life (QoL) are key outcomes in most interventional clinical trials conducted in people living with cystic fibrosis. However, no robust pre-clinical surrogates for FEV1 and QoL exist. The precise physiological mechanisms leading to treatment-related improvements in these outcomes are incompletely understood. In this post-hoc analysis we explored the relationship between changes in the sputum proteome and these outcomes with the aim of identifying translational biomarkers.

**Methods:** Paired sputum samples collected during the AZTEC-CF study (NCT02894684) pre and post 14 days of antibiotic treatment for an acute pulmonary exacerbation were included. Samples were analysed using *in vitro* Mesoscale Discovery (MSD) assays and by nano LC-MS/MS. Peptide identification and quantification was performed and the log-fold change for individual proteins and relationships between protein change and changes in FEV1 and QoL were evaluated.

**Results:** Distinct patterns were found between proteins that correlated with FEV1 and those that correlated with QoL improvements. FEV1 improvement was characterised by increases in bacterially-derived proteins accompanied by decreases in proteins relating to neutrophil degranulation. Conversely, changes in QoL were associated with increases in antiprotease and antioxidant proteins. MSD analysis revealed changes in some neutrophil-associated markers significantly correlated with FEV1 improvements, but no markers significantly correlated with QoL improvements.

**Conclusions:** These results suggest changes in two key CF clinical trial outcomes (FEV1 and QoL) may be underpinned by different physiological mechanisms. Understanding these divergent mechanisms is vital to fortify optimal clinical trial design in CF and panels of biomarkers may be needed to improve translational confidence.

## Introduction

The outlook for people with cystic fibrosis (pwCF) has improved dramatically in the last few decades, thanks in part to the development of highly effective modulator therapies [1]. However, this improved survival may come with an increased lifetime antibiotic exposure, with the potential to develop further antimicrobial-resistant respiratory infections. Furthermore, some pwCF are ineligible for or unable to take modulator treatment [2] and the development of new antimicrobial therapies is therefore needed.

At present, for chronic respiratory infection, there is no robust method to assess the development of new antimicrobial therapies in the preclinical setting that directly aligns with the endpoints used in clinical trials. Many preclinical measures are purely microbiological, and whilst researchers and clinicians can gather information on therapeutics that may effectively control a bacterium experimentally, in the clinical setting complete resolution or clearance of chronic lung infection is rarely achieved. Nevertheless, antibiotics are effective in the treatment of chronic respiratory infections, with improvement in lung function and quality of life [3, 4]. There are currently no preclinical models in which clinical outcomes or their validated surrogates can be measured, and this represents a key hurdle in drug development for CF and the wider chronic lung infection field.

To address these challenges, we investigated potential physiological surrogates of lung function and quality of life changes that could serve as pre-clinical biomarkers for antibiotic drug development models in chronic lung infection.

## Methods

### Study design and datasets

We performed a multi-cohort investigation of the inter-relatedness of changes in lung function and quality of life in response to antibiotics and explored the underlying mechanisms. Firstly, we conducted a post-hoc analysis of existing clinical trial data with external validation, and secondly explored potential biomarkers using banked clinical trial samples.

#### Clinical trial datasets

To investigate the relationship between changes in lung function and in quality of life, we performed a post-hoc analysis of the AZTEC-CF and EvPEx studies [5, 6]. AZTEC-CF (NCT02894684) was a randomised clinical trial conducted in the UK comparing treatment strategies for acute pulmonary exacerbations of infection in pwCF. EvPEx was a prospective observational cohort study assessing lung function dynamics during acute pulmonary exacerbations conducted in Germany (DRKS00012924). The studies have previously been described in detail, but briefly, both included changes in lung function (change in percent predicted forced expiratory volume in 1 second, ppFEV1) and patient reported quality of life (Respiratory domain on the revised Cystic Fibrosis Questionnaire Respiratory domain, CFQ-R). In AZTEC-CF cultured bacterial load (Colony forming units per ml, CFU/ml) data was also available. In each study deidentified individual participant data (IPD) for each outcome was recorded and plotted against each other, with correlation coefficients calculated.

### Sputum biomarker exploration

To quantify putative biomarkers, we utilised banked sputum samples from AZTEC-CF and explored the relationship between changes in clinical outcomes, the sputum proteome and inflammatory profiles. Sputum samples were taken either on Day 1 (pre-initiation of treatment for acute pulmonary exacerbation) or Day 14 (treatment completion). During this trial, samples had been split and either frozen at -80°C, as collected, or treated with sputasol before freezing. Sputasolised samples were used for Mesoscale discovery (MSD) assays and neat sputum samples were prepared for Liquid Chromatography Mass Spectrometry (LC-MS), as described below.

#### Mesoscale Discovery Assays (MSD)

Twelve host-associated inflammatory biomarkers (see Supplementary Table 1) in keeping with guidance from the European Cystic Fibrosis Society Clinical Trial Network Biomarker Working Group [7] from sputum samples were evaluated using the MSD assay platforms (Mesocale Diagnostics, LLC), as per the manufacturer’s instructions. Briefly, sputasolised sputum was centrifuged for 5 minutes at 2000 x *g* to remove bacterial and cell debris, and the supernatant was used to measure biomarker expression. Apart from CRP, for which the assay plates came pre-coated, assay plates were coated with biotinylated capture antibody and incubated at 4°C overnight. Next, plates were washed three times with PBS-Tween. Next, 25 µL of calibrator controls (standards) or samples were added to each well and plates were incubated for 1 hour (2 hours for CRP assay). Plates were washed with PBS-Tween before adding 50 µL (25 µL for CRP assay) of detection antibody to each well and then incubated for a further 1 hour at room temperature. Plates were then washed with PBS-Tween prior to adding Read Buffer to each well and immediately reading outputs using the MESO QuickPlex SQ 1200MM plate reader (Mesoscale Discovery, LLC).

#### Liquid Chromatography Mass Spectrometry analysis

Weighed neat samples were prepared by adding 7 volumes of 80% LC-MS grade Methanol and stored at -80°C until processing. Samples were pelleted by centrifugation at 14000 x*g* before being homogenised using 7M urea buffer. Then, 5 μg protein of each sample was reduced, alkylated, and digested via trypsin in-gel. The resulting peptide mixture was anaylsed via nanoLC-MSMS. Samples were loaded onto the system with a two-minute trapping step before separation on a HSS T3 capillary column (Waters) with a 65 minute gradient (mobile phase A: 0.1 % formic acid; mobile phase B: 0.1 % formic acid in acetonitrile). Samples were introduced to the mass spectrometer (Exploris 240, Thermo Fisher) via electrospray ionisation and analysed in data dependent acquisition mode. Raw data was processed using Progenesis QIP 4.2 (Nonlinear Dynamics) for quantification and Thermo Proteome Discoverer 2.5 (Thermo Fisher) for identification purposes. Respective swissprot databases (human, *Pseudomonas aeruginosa, Staphylococcus aureus, Veillonella parvula, and Veillonella dispar)* were obtained from uniprot.org. These bacterial species were chosen as the most abundant species in 16S rRNA sequencing of the samples published previously and based on available high-quality databases.

### Statistics

Analyses were conducted in Rstudio, and data are presented as median (IQR) or mean (SD) throughout depending on normality. Associations between outcomes were evaluated using Pearson’s correlation coefficient.

## Results

### 1/ Individual participant data analyses reveal poor correlation between changes in lung function and quality of life in prospective clinical trials

Participant characteristics for the AZTEC-CF and EvPex studies are previously described but in brief, median (IQR) age was 29.5 yrs and 28.5 yrs, and median FEV1 was 52% predicted and 43% predicted respectively.

Overall lung function improvements were similar between studies with a +9.5% improvement in lung function seen over the course of the EvPex study and +11.8% improvement in lung function seen in AZTEC-CF. Similarly, CFQ-R respiratory domain increased +17 points in EvPex and +12.5 points in AZTEC-CF.

Individual lung function responses to each study are presented in waterfalls plots in Figure 1A and 1B with each participant’s lung function response colour coded in order of magnitude. Individual changes in QoL are also presented in Figures 1C and 1D, again with the colour codings signifying lung function response. These waterfalls plots demonstrate heterogeneity in lung function and QoL responses in both studies with some of the poorest lung function responders achieving excellent QoL improvements and vice-versa. Similarly, correlation between changes in lung function and quality of life in both studies was poor (r=0.25 and r=0.29 respectively for AZTEC-CF and EvPEx, Figure 2).

**Figure 1:**
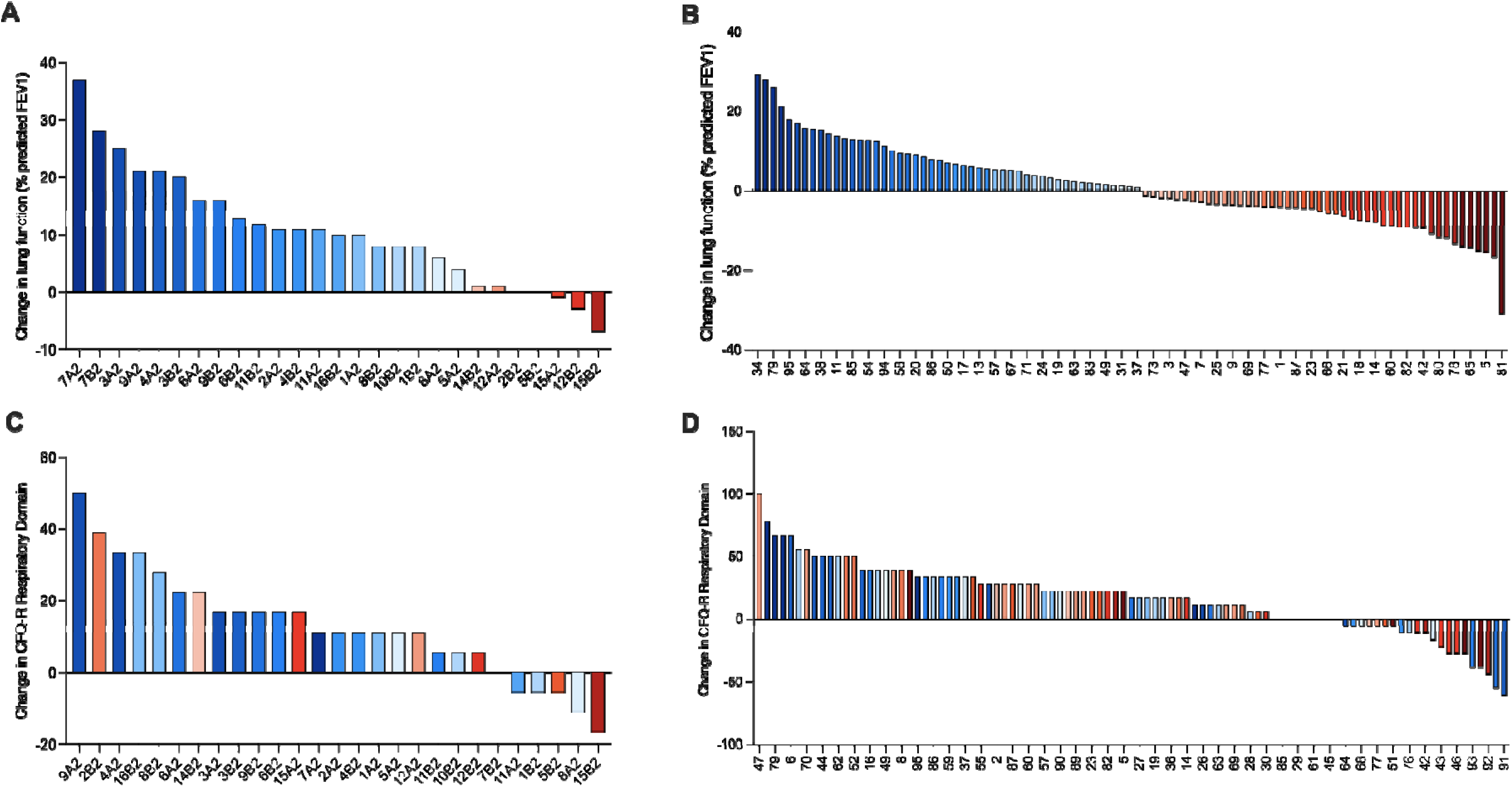
Changes in FEV1 and QoL scores by individual participant during a clinical trial. The relative changes in FEV1 and QoL scores (measured using CFQ-R) collected per participant of the AZTEC-CF and the EvPex studies show weak correlations between these measures. A) Change in lung function (% predicted FEV1) coloured by order of magnitude from most improved to least improved for participants in the AZTEC-CF study. Study participants (Numerical ID) were shown to have differing responses at either intervention A or intervention B. B) Change in lung function(% predicted FEV1) for all participants of the EvPex study ordered and coloured according to the magnitude of change from most improved to least improved. C) Change in CFQ-R Respiratory domain for AZTEC-CF participants organised in order of magnitude from most improved to least improved with colour codes signifying lung function response. D) Change in CFQ-R Respiratory domain for EvPex participants organised in order of magnitude from most improved to least improved again with colour codes signifying lung function response. For some participants there was a clear correlation between improved scores in FEV1 and QoL whereas for others improvements in one score correlated with worsening of the other score.

**Figure 2:**
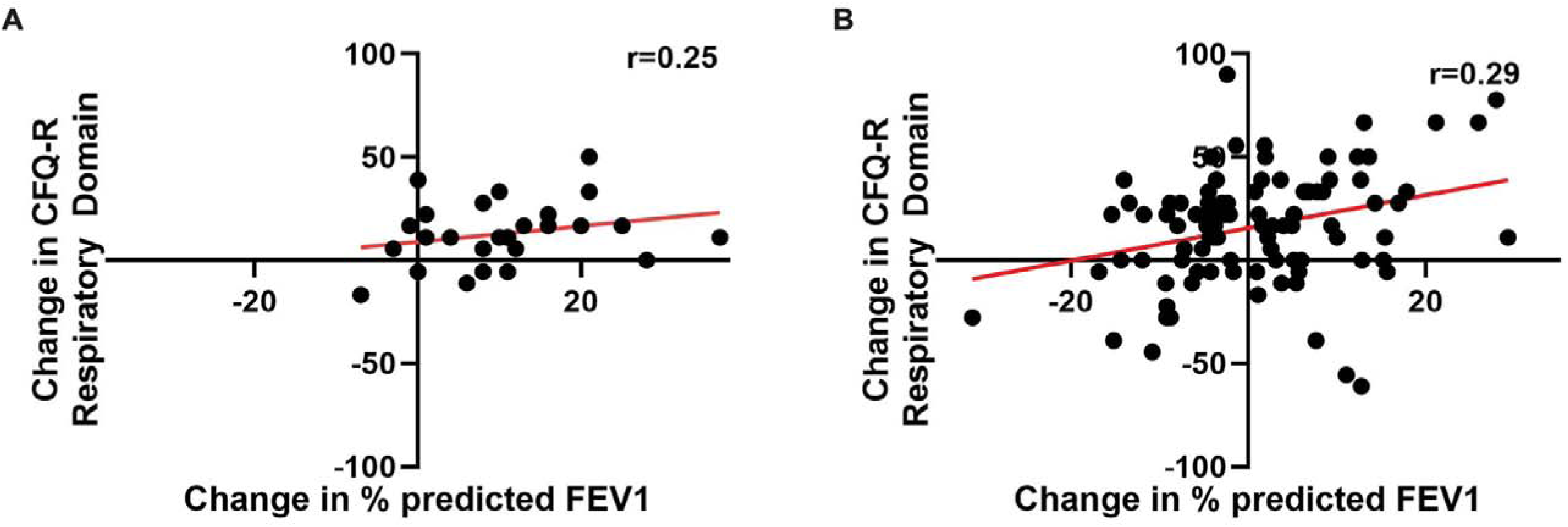
Weak correlations between lung function and Quality of Life changes during interventional clinical trials for pwCF. A) Weak associations between lung function changes with changes in CFQ-R Respiratory Domain for participants in the AZTEC-CF study. B) Similar weak associations between lung function changes with changes in CFQ-R Respiratory Domain for participants in the EvPex study when analysed using Spearman’s correlation analysis.

In AZTEC-CF, cultured bacterial load data were available and this again showed poor correlation with improvements in either lung function or QoL, (Supplemental Figure 1).

### 2/ Changes in quality of life and lung function are associated with distinct changes in the untargeted sputum proteome

To explore whether the observed divergent responses to lung function and quality of life are associated with differential biological processes we performed untargeted proteomics in banked sputum samples from the AZTEC-CF study. Paired sputum samples from the same individual pre- and post-treatment were used to determine key signatures or markers associated with changes in clinical outcomes (FEV1 or QoL). Protein identification was based on at least two unique peptides. This resulted in 712 proteins being identified (Repository accession number?). We used linked metadata between sputa and clinical trial outcomes to derive correlations between the fold change in abundance for individual proteins and changes in both FEV1 and QoL. These analyses revealed little apparent overlap between proteins that were strongly associated with changes in FEV1 and those with changes in QoL (Figure 3A). Heatmap analysis showed that significant increases or decreases in the abundance of proteins correlated with FEV1 scores had weaker or opposing correlations when compared to QoL scores. This finding was confirmed at the individual protein level, where proteins that were significantly associated with changes in FEV1 were completely distinct from those associated with QoL (Figure 3B and 3C).

**Figure 3:**
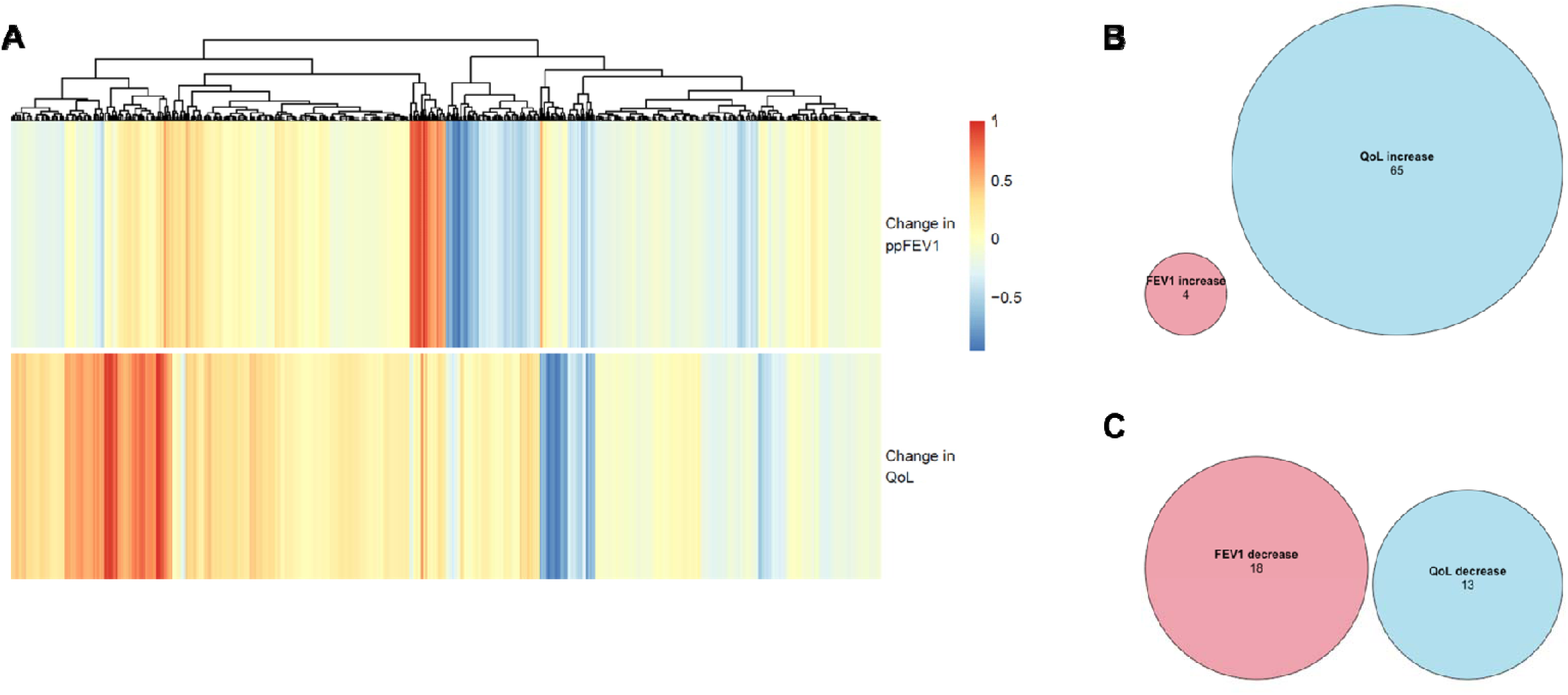
Protein abundance changes related to FEV1 or QoL show distinct signatures. A) Heatmap showing that changes in the abundance of proteins detected in the sputum of pwCF are differential when comparing this to changes in either FEV1 or QoL outcomes. B) Euler diagram showing there was no overlap in proteins that were significantly increased in abundance when FEV1 or QoL outcome measures improved. C) Euler diagram showing there was no overlap in proteins that were significantly decreased in abundance when FEV1 or QoL outcome measures improved

Furthermore, inspection of the top 30 differentially abundant proteins revealed distinct pathway associations for FEV1 and QoL changes (Figure 4A and Figure 5A).

**Figure 4:**
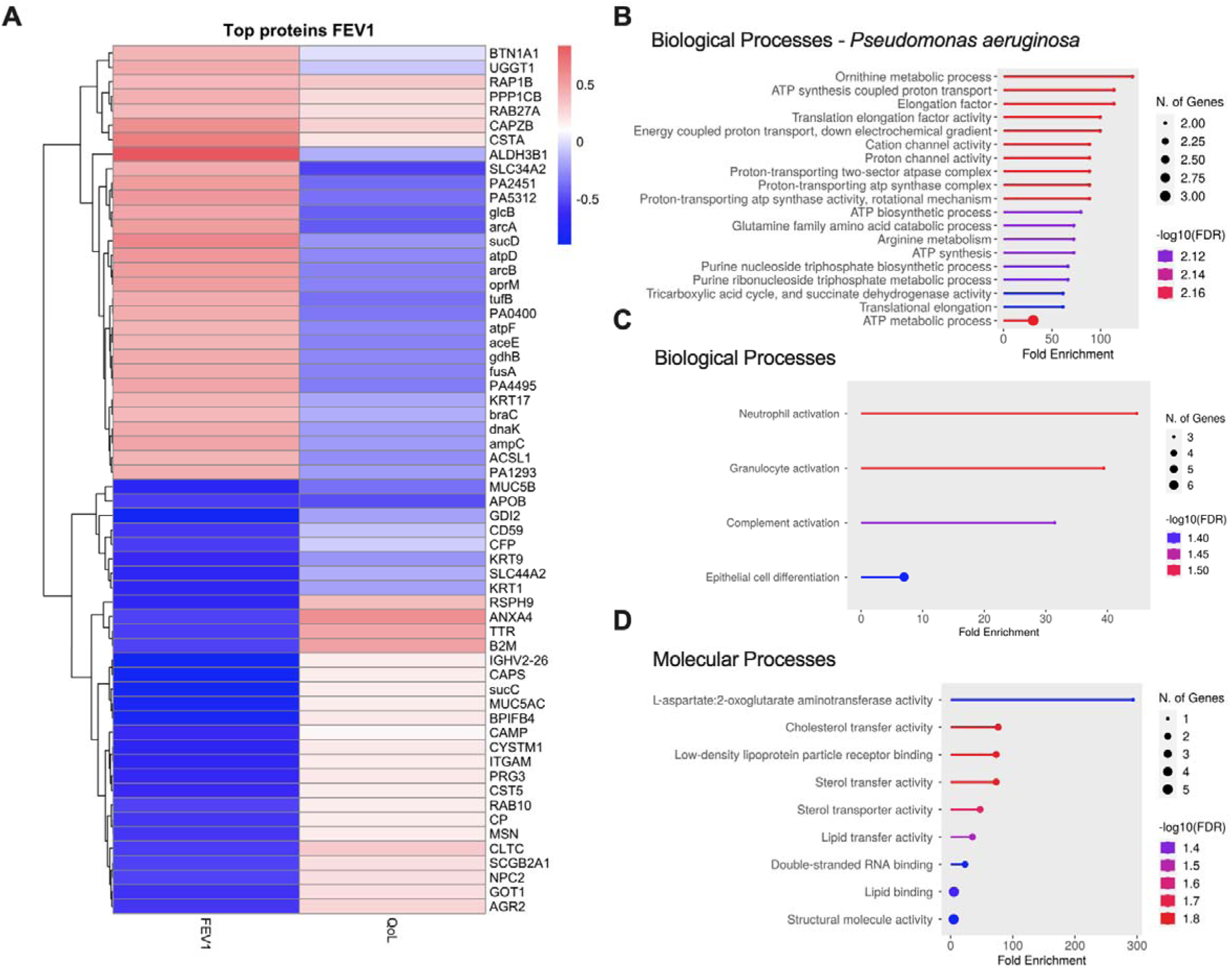
Hierarchical clustering and pathway prediction for the top proteins that were differentially increased or decreased when FEV1 improved. A) The top proteins that increased (n=30) and decreased (n=30) when FEV1 improved following exacerbation resolution in the AZTEC-CF trial determined based on lowest p-values. B) Biological processes and C) Molecular processes most associated with the human proteins that were downregulated in correlation with FEV1 improvements were determined using the ShinyGO server (http://bioinformatics.sdstate.edu/go/) and ranked by fold enrichment.

**Figure 5:**
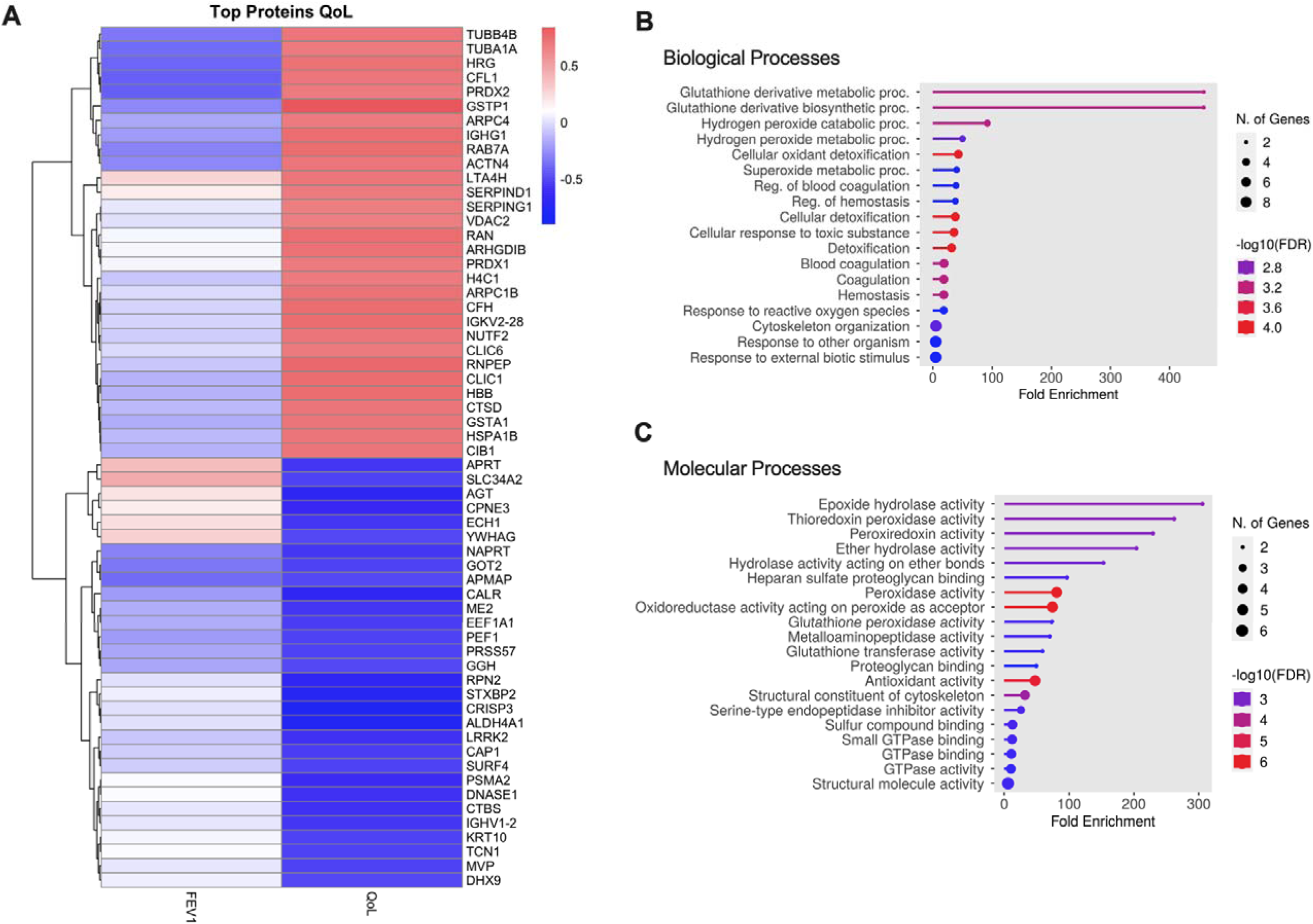
Hierarchical clustering and pathway prediction for the top proteins that were differentially increased or decreased when QoL improved. A) The top proteins that increased (n=30) and decreased (n=30) when QoL improved following exacerbation resolution in the AZTEC-CF trial determined based on lowest p-values. B) Biological processes and C) Molecular processes most associated with the human proteins that were upregulated in correlation with QoL improvements were determined using the ShinyGO server (http://bioinformatics.sdstate.edu/go/) and ranked by fold enrichment.

Focusing on the top 30 proteins that differentially increased in abundance with FEV1 improvements, 20/30 were bacterial proteins associated with the pathogen *Pseudomonas aeruginosa* (Figure 4B). The remaining 10 proteins were host-associated but did not appear to have any clearly associated host physiological processes when input into the STRING protein-protein association networks database (Szklarczyk et al., 2023, Supplemental Figure S2A) or when using the ShinyGO server to analyse protein Gene Ontology (http://bioinformatics.sdstate.edu/go/). Analysis of the top 30 proteins that differentially decreased in abundance when FEV1 improved showed that 29/30 of these were host-associated. Analysis of Gene Ontology for these proteins highlighted a reduction in neutrophil and granulocyte activation accompanied by a reduction in proteins associated with complement activation (Figure 4B). There was also a reduction in the protein levels for MUC5AC, MUC5B, and AGR2, all notably associated with salivary secretions [8] (Supplemental Figure 3B).

Of the top 30 proteins differentially associated with QoL improvements, 0/30 were bacterially-derived (Figure 5A). All these host-derived proteins were completely distinct from those that were significantly associated with FEV1. Key proteins that were upregulated when QoL improved are involved in antioxidant or antiprotease activity. In particular, there were a large number of proteins found to be associated with processes involved in reduction of peroxidases or sequestering free-radicals, which can cause damage in the airways. In contrast, proteins that were differentially decreased in abundance in association with QoL improvements were observed to be mostly cellular components such as lysosymes, granules, vacuoles and organelles, with no specific biological or molecular processes associated (Supplemental Figure 4).

### 3/ No individual inflammatory markers are associated with both lung function and quality of life outcomes

To test the relationship between changes in clinical trial outcomes and targeted inflammatory mediators, the same sputum samples were tested against a panel of 12 host-associated biomarkers (Supplementary Table 1). Of these, Calprotectin and IL-12p70 were undetectable in all samples tested. The quantity of each of the remaining ten biomarkers was measured in samples pre- and post-treatment. Changes in the levels of biomarkers between Days 1 and Day-14 were then analysed in comparison with the corresponding change in FEV1 or QoL score, using Spearman’s correlation analysis.

This analysis showed that reductions in the levels of 5 host-associated biomarkers (MPO, MMP-9, VEGFA, YKL-40 and IL1-β) were found to have significant correlations with improvements in FEV1 score however, no biomarkers were significantly associated with changes in QoL scores. The correlations were much weaker or were directionally opposed to the correlations observed with FEV1 scores (Supplemental Figure 5 and 6). Heatmap visualisation of these correlations clearly depicts these differences (Figure 6). MMP-9 has the closest correlations with both FEV1 (r = 0.50, *p* = 0.017) and QoL though this was not significantly associated for QoL (r = 0.36, *p* = 0.099) (Figure 6 and Supplemental Figure7).

**Figure 6:**
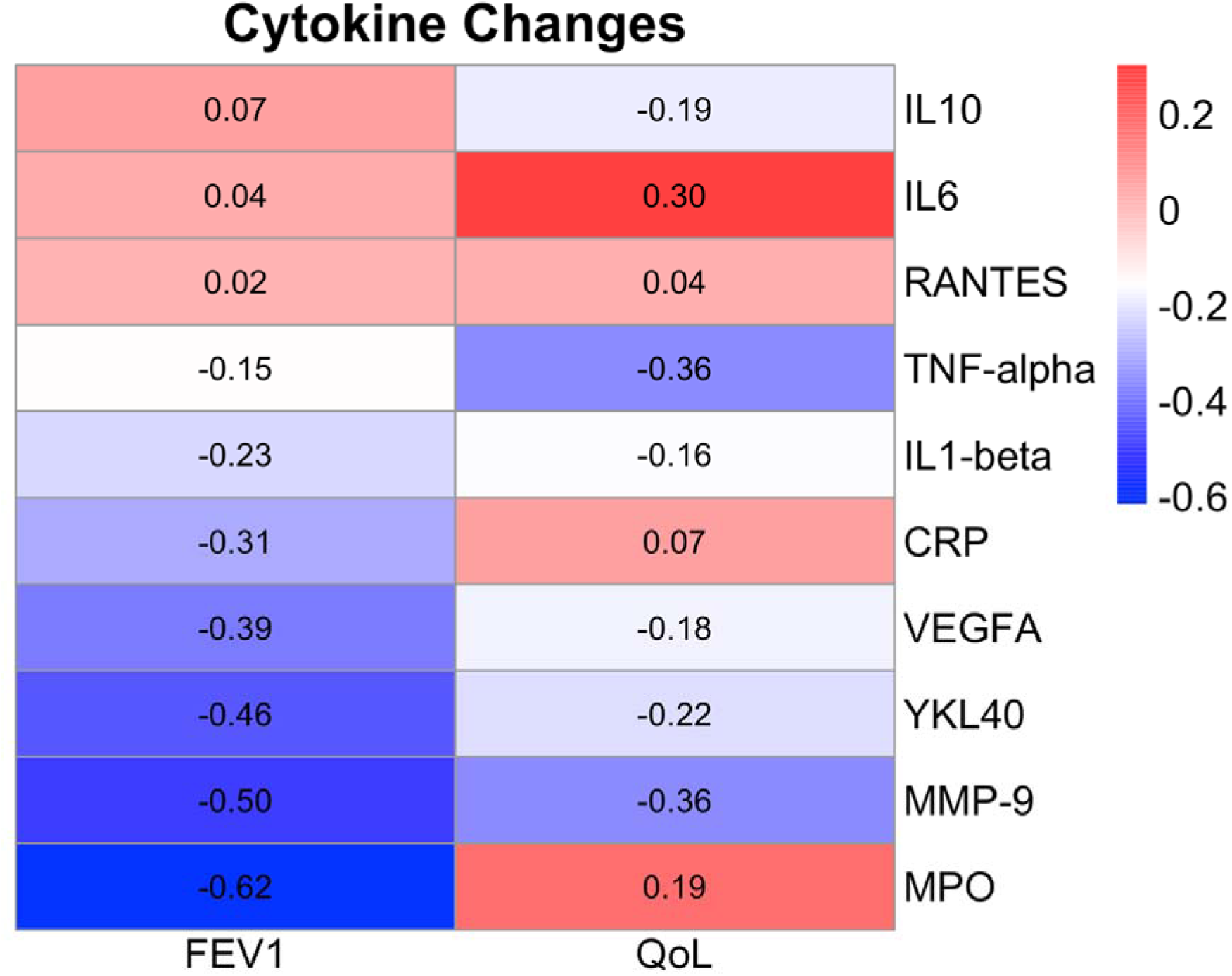
Heatmap of Spearman’s correlation scores for host-associated biomarker alterations in Sputum from pwCF. Sputum from participants of the AZTEC-CF trial was screened using a panel of 10 host-associated biomarkers. Spearman’s correlation analysis was used to assess associations between the change in sputum levels at day-14 and the change in either FEV1 or QoL at day-14. For MPO, MMP-9, YKL40 and VEGFA were the only biomarkers to have significant associations with FEV1 improvements. No biomarkers showed significant associations with QoL improvements. MMP-9 was the only biomarker to have strong associations in the same direction for both FEV1 and QoL, though for QoL this was not significant.

## Discussion

In this study we investigated the relationship between changes in FEV1 and QoL in two CF cohorts and explored potential biomarkers that could facilitate pre-clinical evaluation of drugs in the chronic lung infection setting. Our main findings are that changes in lung function and QoL are associated with distinct host-pathogen processes with differing neutrophil-associated roles being key to each outcome (Figure 7). The major implication is that a single biomarker will not accelerate translational confidence in pre-clinical chronic lung infection models, and instead composite biomarker panels may be required.

**Figure 7:**
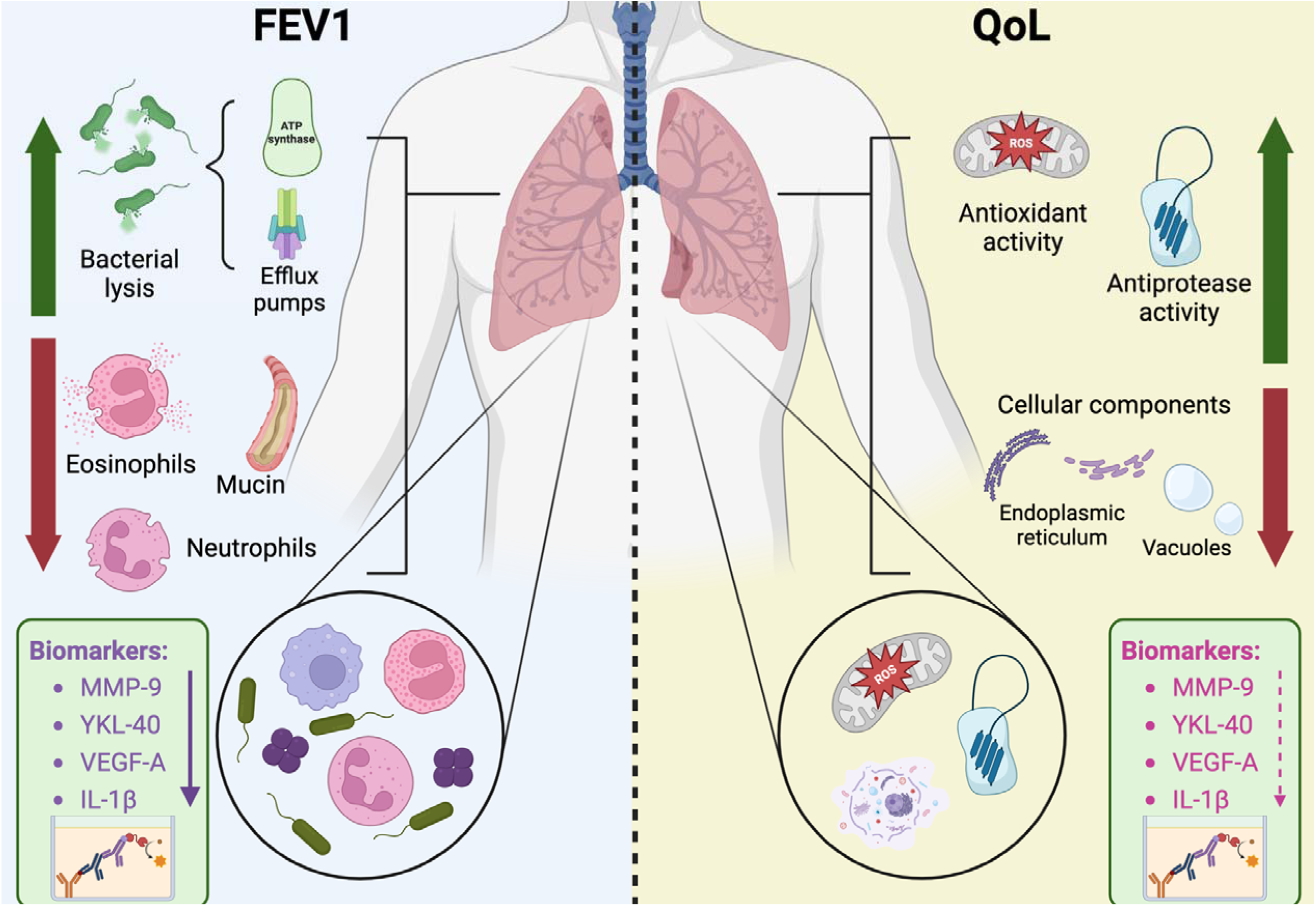
Host and pathogen physiological changes associated with CF clinical trial outcomes. A schematic diagram for the key host and pathogen physiological pathways determined to be associated with improvements in either FEV1 or QoL following treatment for an acute pulmonary exacerbation. Neutrophil-driven responses were clearly associated with both FEV1 and QoL improvements, however the differential proteomes associated with each outcome measure revealed that different neutrophil-associated roles are at play. FEV1 improvements are defined by increases in bacterial proteins suggestive of bacterial release from biofilms associated with chronic lung infections. Neutrophil degranulation and NETosis-related proteins were decreased in association with FEV1 improvements. Conversely, QoL is markedly defined by increases in antioxidant and antiprotease proteins, potentially indicative of the restoration of imbalances in these.

Lung function and QoL are key outcomes in most interventional clinical trials of antimicrobial therapeutics conducted in pwCF. Here, we analysed data from two clinical trials (one RCT and one observational study) in different countries encompassing different exacerbation settings (hospital only for AZTEC-CF, home/hospital for EvPEx). Despite differences in the design of each study, we consistently found only weak correlations between changes in FEV1 and QoL in response to treatment. The relationship between outcomes variables in clinical trials is rarely reported, for example a meta-analysis of RCTs published in high-impact journals estimated ∼1% of studies reported the relationship between outcomes [9]. However, understanding the relationship between outcomes can inform investigators with respect to causal mechanisms and pathways as well as informing future study design. This is particularly relevant to CF where despite the routine use of FEV1 and QoL in studies of antimicrobials, the precise pathophysiological changes associated with each outcome measure and how these respond to different treatments are poorly understood. Moreover, FEV1 and QoL are impossible to measure pre-clinically and there are no well validated surrogates. Bacterial load is often therefore relied upon during early-stage drug development, however, the traditional paradigm of antimicrobials reducing bacterial load and thereby leading to improved lung function has been consistently disproved. For example, multiple studies in the chronic lung infection setting have observed clinical improvements upon treatment that are not associated with measurable changes in infection burden [3, 4]. Illustratively, in AZTEC-CF, bacterial load did not correlate with changes in lung function or QoL. The challenges inherent in making drug efficacy predictions in the absence of quantifiable preclinical endpoints that feed through into the clinical setting represent a significant barrier to effective translational drug development and efforts are needed to overcome them.

Proteomics revealed striking differences in the proteins associated with changes in lung function and QoL. For example, across the entire proteome, there was no similarity or crossover in the proteins for which abundance changes significantly associated with changes in FEV1 or QoL, suggesting different underlying biological mechanisms at play. This was supported by pathway analyses which found neutrophil and granulocyte activation were key pathways associated with change in lung function, whereas glutathione regulatory processes were linked to changes in QoL. Exaggerated neutrophilic airway inflammation is well characterised in CF and changes in neutrophil-derived protein abundance have been associated with lung function cross-sectionally. [10–13] Our observation of decreased neutrophil-derived protein abundance as a function of FEV1 improvements, following treatment, supports these earlier studies but goes further in suggesting neutrophil activity is at least a biomarker of changes in lung function and could be a putative therapeutic target.

The main proteins accountable for the relationship between neutrophil-derived proteins and lung function were the antimicrobial peptide cathelicidin (CAMP), proteoglycan 3 (PRG3) and integrin alpha M (ITGAM). CAMP and PRG3 are associated with the release of Neutrophil Extracellular Traps (NETs), an increase of which has been described in CF sputum [12, 14, 15]. Whilst the major role of NETosis during infection is to trap and kill bacteria, the release of other neutrophil components such as DNA contributes to the viscous airway secretions that cause airway obstructions that could result in reduced FEV1, thus providing a putative mechanistic link between the proteomics change we observed and changes in lung function [16].

Similarly, MUC5AC, MUC5B and AGR2, all involved in mucin production, were also decreased in correlation with FEV1 improvements. MUC5AC and MUC5B are two major gel-forming mucins that are commonly detected in CF airways and may also contribute to mucus plugging of the airways with subsequent obstruction and reduction in FEV1. The decreases in abundance of MUC5AC and MUC5B observed in this study are complementary to results found in previous studies where they were the predominant mucins detectable in CF sputum samples [17–19]. Henke et al. [19] showed that both mucins were increased in CF airway secretions during exacerbation and, similar to Voynow et al. [20], they hypothesised that this increase during exacerbation was likely a protective mechanism. Here, MUC5AC and MUC5B were found to decrease in abundance when FEV1 improved, presumably due to exacerbation resolution following treatment, when their protective response was no longer required. These mucins could therefore also prove useful as *in vitro* biomarkers.

Key pathways involving the top 30 upregulated proteins associated with QoL improvements were antioxidant/oxidant detoxification pathways and antiprotease/protease inhibition pathways. For pwCF, there is an imbalance in both proteases/antiproteases and oxidants/antioxidants that contribute to CF pathophysiology [21–23]. The main contributors to the release of proteases and oxidants in the CF lung are the infiltrating neutrophils, which release high levels of neutrophil elastase (NE) [24], MMP-9 [25] and oxidants such as MPO [26]. Maher *et al.,* [12] showed that these imbalances get progressively worse with disease severity. We have shown that treatment for acute pulmonary exacerbation results in increases in serine-type antiprotease-related proteins that could be associated with reduced protease activity culminating in the relief of inflammatory symptoms.

Sloane *et al.,* [27] highlighted the significant effect that MPO has on pulmonary inflammation in CF lungs. In addition, lower levels of antioxidant proteins such as glutathione-S transferase (GST) and peroxiredoxin were detected in CF sputum compared to healthy controls [28–30]. In this study, proteins associated with GST (GSTA1 and GSTP1) and peroxiredoxin (PRDX1 and PRDX2) were significantly increased in abundance when QoL improved. These increases could be associated with better control of the oxidants (MPO) that are released by neutrophils during NETosis, following recruitment to the lungs. Combined, the increases in both antiprotease and antioxidant proteins when QoL improved could be linked to better control of neutrophilic inflammation following exacerbation resolution in pwCF. As these changes were uniquely associated with QoL improvements and not FEV1 improvements, it could prove beneficial for clinical studies looking at the effect of antiprotease/antioxidant treatment in pwCF to utilise QoL outcome measures as superior to FEV1 when assessing the success or failure of such therapies. For example, glutathione has long been thought to play a role in the development and progression of CF lung disease [31]. Nebulised glutathione supplementation has previously been investigated for pwCF [32]. Interestingly, those trials, powered towards lung function improvement, were negative, but glutathione has been found to improve QoL in people living with chronic pancreatitis [33] and our data could suggest QoL may be a more suitable trial endpoint.

Bacterial proteins dominated the top proteins associated with improvements in lung function. Out of the 20 bacterial proteins highly correlated with improved FEV1 measures, 6 (30%) were shown to be among the most highly expressed and conserved genes within *P. aeruginosa* genomes [34]. Collated data on gene expression on the *Pseudomonas* genome database [35] also found them to be among the most highly expressed in CF sputum. Many of the bacterial proteins detected were highly abundant proteins associated with ATP production or energy metabolism which could suggest the presence of bacterial cells within the sputum. These increased proteins could be indicative of the release of bacterial cells from embedded biofilms within the lungs following antimicrobial treatment and this could also explain the increased abundance of total bacterial and *P. aeruginosa* load found in the sputum in this study and the lack of overall correlation between sputum bacterial load and clinical endpoints in general. Overall, we found a lack of crossover in proteins associated with either FEV1 or QoL improvements. This suggests that a panel of biomarkers may be required to efficiently determine potential clinical trial success during pre-clinical testing of novel therapeutics.

To understand if readily available commercial biomarkers could form the basis of such a panel we evaluated 10 biomarkers recommended by the ECFS biomarker group [7]. We were able to show significant treatment-associated reductions in the sputum levels of MPO, MMP-9, YKL-40, VEGF-A and IL-1β, all of which are known to be increased during pulmonary exacerbations [36–40]. However, although we found promising results for some biomarkers as surrogates for FEV1 improvement, no biomarkers were significantly associated with QoL. These findings further reinforce our earlier findings of distinct drivers of change for the respective endpoints but also suggest bespoke panels will likely be needed if preclinical models wish to effectively predict clinical improvements. We confirmed that changes in neutrophil-derived proteins can be detected using simple *in vitro* assays. MMP-9, a metalloprotease detected in high levels in CF sputum, was the only biomarker to show similar correlations with both FEV1 and QoL improvements and warrants further investigation into its potential as part of a biomarker panel.

This study has several limitations. The proteomics and cytokine assays were performed on sputum samples from a small number of pwCF who had taken part in an antimicrobial intervention study at a single centre in the UK. Larger scale studies should be performed to ensure that findings are translatable across cohorts. Similarly, the studies herein use CFQ-R to characterise changes in quality of life, validation across other HR-QOL tools is required. Additionally, sputum samples used in this study were taken before modulator therapies were licensed for use in the UK and therefore the effect modulators have on exacerbation resolution following antimicrobial intervention remains unknown. A recent study by Maher et al (2023) [41] showed that modulator therapy had similar effects on the sputum proteome as shown here, however, this did not lead to a fully healthy lung proteome but rather a more intermediate state still indicative of altered lung inflammation. It therefore remains to be seen what effect modulators may have on physiological responses in pwCF following pulmonary exacerbations and the relationship with potential biomarkers.

In conclusion, this study demonstrated that the key clinical trial outcome measures FEV1 and QoL are associated with different physiological processes in pwCF. These findings suggest the need for bespoke panels of translational biomarkers and further work is needed to prospectively identify, optimise and validate these approaches to allow improved confidence and efficiency in future drug development.

## Acknowledgements

JF and DN would like to acknowledge funding from the CF Trust and the CF Foundation (SRC022).

