## Supplemental Material for "Sputum Proteomics reveals unique signatures linked to key outcomes in cystic fibrosis trials"

**Supplementary Material**

**Clinical cohort information**

We performed analyses for this study using data from two independent cohorts from the AZTEC-CF and EvPEx studies respectively. The studies were approved by the relevant national research ethics and regulatory committees. All patients provided informed consent and approval numbers were 16/NW/0741 and DRKS00012924. The studies both have been previously been described and results have been published but are also described below:

**AZTEC-CF:**

Aztreonam lysine for the treatment of exacerbations of cystic fibrosis (AZTEC-CF):

This was an open-label randomised cross-over trial comparing two treatment strategies for acute exacerbations of cystic fibrosis. The study was conducted at the adult CF centre at Liverpool Heart & Chest Hospital NHS Foundation Trust.

Key inclusion criteria were age >= 16yrs of age, clinical diagnosis of cystic fibrosis, percent predicted FEV1 between 25 and 75%, Chronic *P. aeruginosa* infection, acute pulmonary exacerbation requiring inpatient intravenous antibiotics. Key exclusion criteria: Prior positive sputum culture for an organism in the *Burkholderia cepacia* complex. Interventions: ecruited pwCF were randomised to receive two treatment regimens sequentially over the course of their next two exacerbations requiring intravenous antibiotics. The threshold for treatment inclusion was standardised using the modified Fuch's criteria, which included the need for intravenous antibiotic treatment and a recent change of least two of the following criteria: change in sputum volume/colour, increased cough, increased fatigue, malaise/lethargy, anorexia/weight loss. Unstratified, block randomisation in groups of four was carried out prospectively using Statsdirect® (Statsdirect Ltd, 2013) and kept in sealed envelopes to be referred to by the investigators at the time of participants’ first exacerbation. Treatment regimens were “AZLI+IV” (14 days of AZLI 75mg tds plus IV colistimethate 2 Mega Units tds) and “IV+IV” (14 days of IV colistimethate 2 Mega Units tds plus a second IV antipseudomonal antibiotic selected by the admitting physician). Primary outcome was change in % predicted FEV1 at day 14. Secondary outcomes included changes in the Respiratory Domain of the CFQ-R health-related quality of life questionnaire and change in cultured bacterial load. Sputum samples were taken prior to start of treatment and at day 14 with specific consent for use in future research. In total 16 participants were enrolled and 28 exacerbation treatments were successfully completed. Paired samples were available for

**EvPEx:**

This nonrandomized, prospective single-centre observational trial (DRKS00012924) was conducted at Cystic Fibrosis Center, Clinic Westbrandenburg, Potsdam, and Charité-Universitaetsmedizin Berlin, Germany. PwCF with acute exacerbation who were started on antibiotic therapy either as outpatients or inpatients were recruited for the study on the day of their presentation. Therapeutic decisions prior to and during the study were made by clinicians regardless of inclusion in the study. The selection criteria for including pwCF were as follows: No history of lung transplantation; ≥6 years or ≤75 years; acute exacerbation as defined by modified Fuch’s criteria. Ability to perform lung function and home spirometry. Exclusion criteria: History of lung transplantation; <6 years or >75 years; non-ability to perform lung function and home spirometry; Outcomes of interest included CFQ-R and lung function. C-reactive protein, sputum microbiology, and body mass index were the additional parameters that were optionally collected. In addition to departmental spirometry at baseline and day 28, follow-up examinations with home spirometry and collection of the symptom questionnaire were carried out at home on days 7, 14, and 21. All parameters were collected again during a follow-up examination on site on day 28. In total 105 participants have been recruited for the study of whom 96 completed the study and of 95 participants paired lung function and HRQOL data could be evaluated.

**Table 1**: List of host-associated biomarkers used during this study.

| **Name of Biomarker** | **Abbreviation** | **Relation to CF or CF exacerbations** | **References** |
| --- | --- | --- | --- |
| Myeloperoxidase | MPO | Present in CF sputum, contributor to parenchymal destruction | Mohammed et al. (1988),Koller et al. (1995), Thomson et al. (2010) & Hair et al. (2017) |
| Matrix Metalloprotenaise-9 | MMP-9 | Present at high levels in CF lower airway secretions, correlated with FEV1 in paediatric pwCF. | Ratjen et al. (2002), Sagel et al. (2005) & Thomson et al. (2010) |
| Chitinase-3 like 1 gene | YKL-40 | Higher levels in serum linked to more severe clinical phenotypes. | Leonardi et al. (2016) & Coriati et al. (2021) |
| Vascular Endothelial Growth Factor | VEGF-A | Upregulated in CF airways and *cftr -/-* mice, involved in vascular remodelling of the airways | Clémence et al. (2013) |
| Interleukin-1β | IL-1β | Detectable in BAL from pwCF and increased during exacerbation | Montgomery et al. (2018) |
| Interleukin-6 | IL-6 | Sputum concentrations correlated positively with FEV1 | Nixon et al. (1998) |
| Interleukin-10 | IL-10 | Involved in control of inflammatory T-cell responses to bacterial infections | Casaulta et al. (2003) |
| Regulated upon Activation, Normal T Cell Expressed and Presumably Secreted | RANTES | Expression is altered in CF epithelia and is upregulated in nasal polyps | Schwiebert et al. (1999) & Scapa et al. (2011) |
| C-Reactive Protein | CRP | Serum level changes commonly used as reporter during exacerbation. Shown to predict treatment non-response and time until next exacerbation | Sharma et al. (2017) |
| Tumour Necrosis Factor alpha | TNF-α | High levels in airways contribute to chronic neutrophil influx and airway damage. | Mitola et al. (2008) |

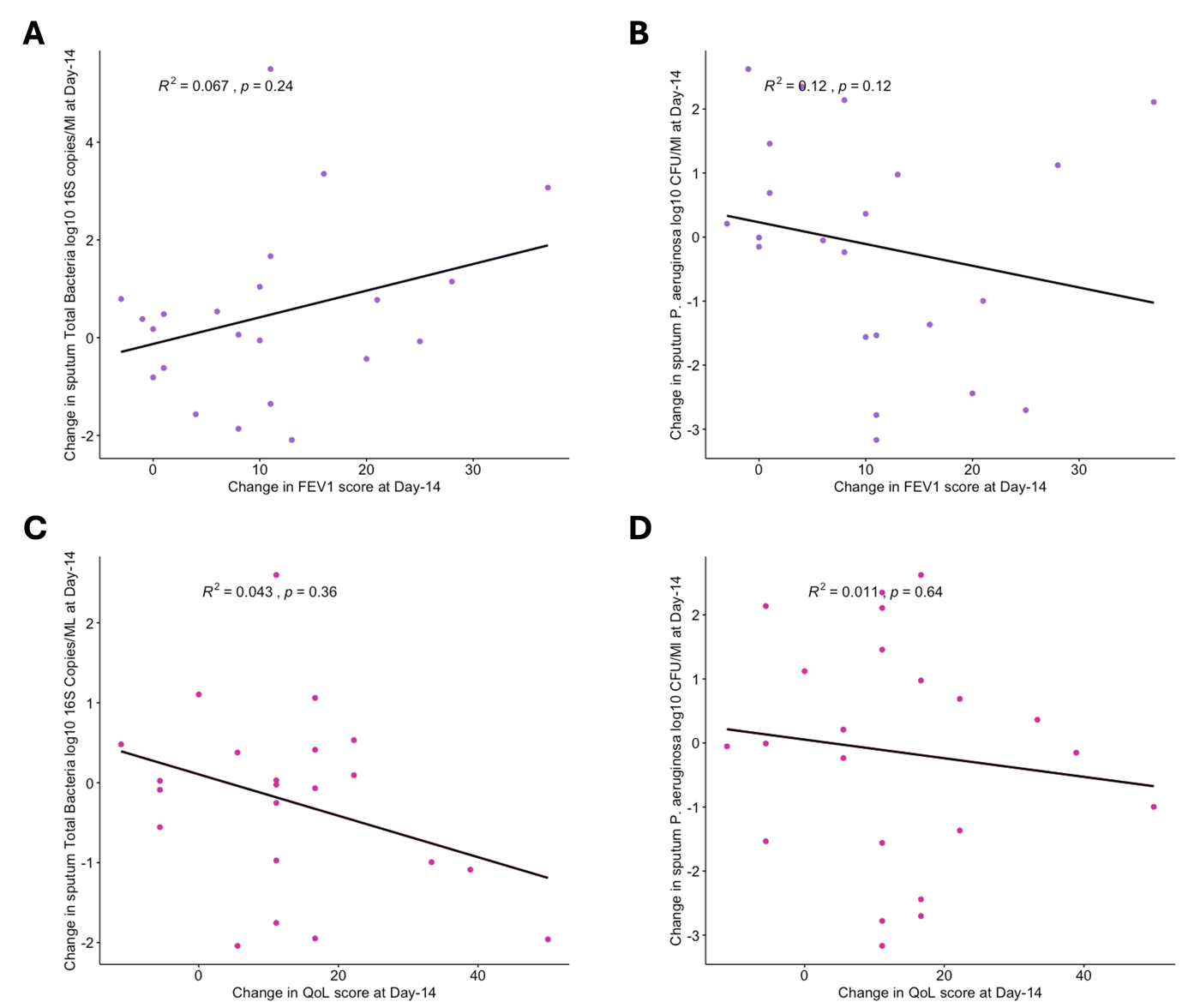

**Supplementary Figure 1: Spearmans correlations for changes in clinical outcomes and bacterial load at Day-14 post antimicrobial intervention.** A) Weak positive correlations between FEV1 improvements at Day-14 total bacterial load as determined by 16S copy number (B) Weak negative correlations between FEV1 improvements and changes in log10 CFU/ml *P. aeruginosa* at day-14 Conversely there were weak negative corelations with both total bacterial load (C) and log10 CFU/ml *P. aeruginosa* (D) and QoL improvements

.

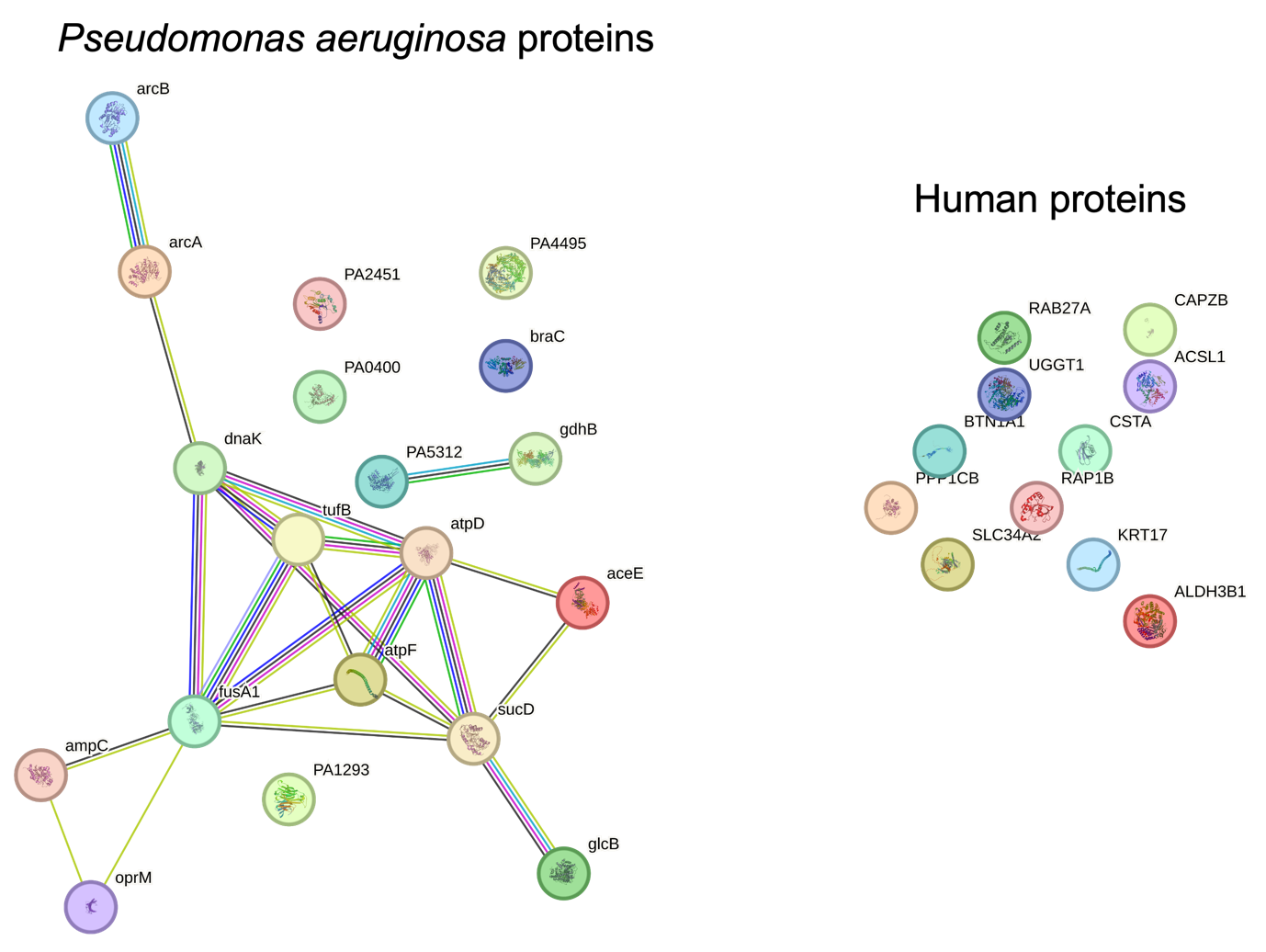

**Supplementary Figure 2: Pathways and Protein-Protein interaction Networks for the top 30 proteins increased in abundance when FEV1 improved.** The STRING database showed that 17/30 proteins were predicted to have some form of interaction based on known interactions that were either experimentally determined or from curated databases, predicted interactions from gene neighbourhood, gene fusions or gene co-occurance and from other sources such as textmining, co-expression or protein homology. Key interactions predicted by STRING for the bacaterial and host proteins ranked in the top30 proteins increased when FEV1 improved. No interactions predicted by STRING for the host proteins ranked in the top 30 proteins increased when FEV1 improved.

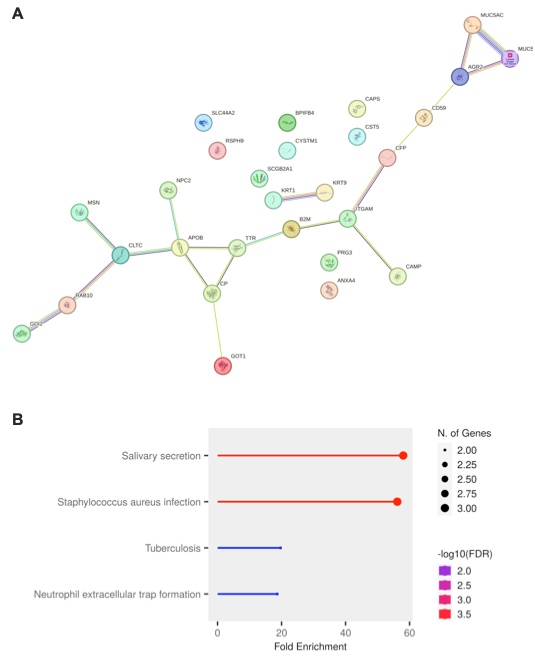

**Supplementary Figure 3: The Protein-Protein interaction Network for the top 30 proteins decreased in abundance when FEV1 improved.** A) The STRING database showed that 17/30 proteins were predicted to have some form of interaction based on known interactions that were either experimentally determined or from curated databases, predicted interactions from gene neighbourhood, gene fusions or gene co-occurance and from other sources such as textmining, co-expression or protein homology. Of this key interaction network many of the proteins appear to be involved in pathogen control or clearance processes, in particular the presence of MUC5AC, MUC5B and AGR2 involved in the production of key mucins detected in CF sputum. B)

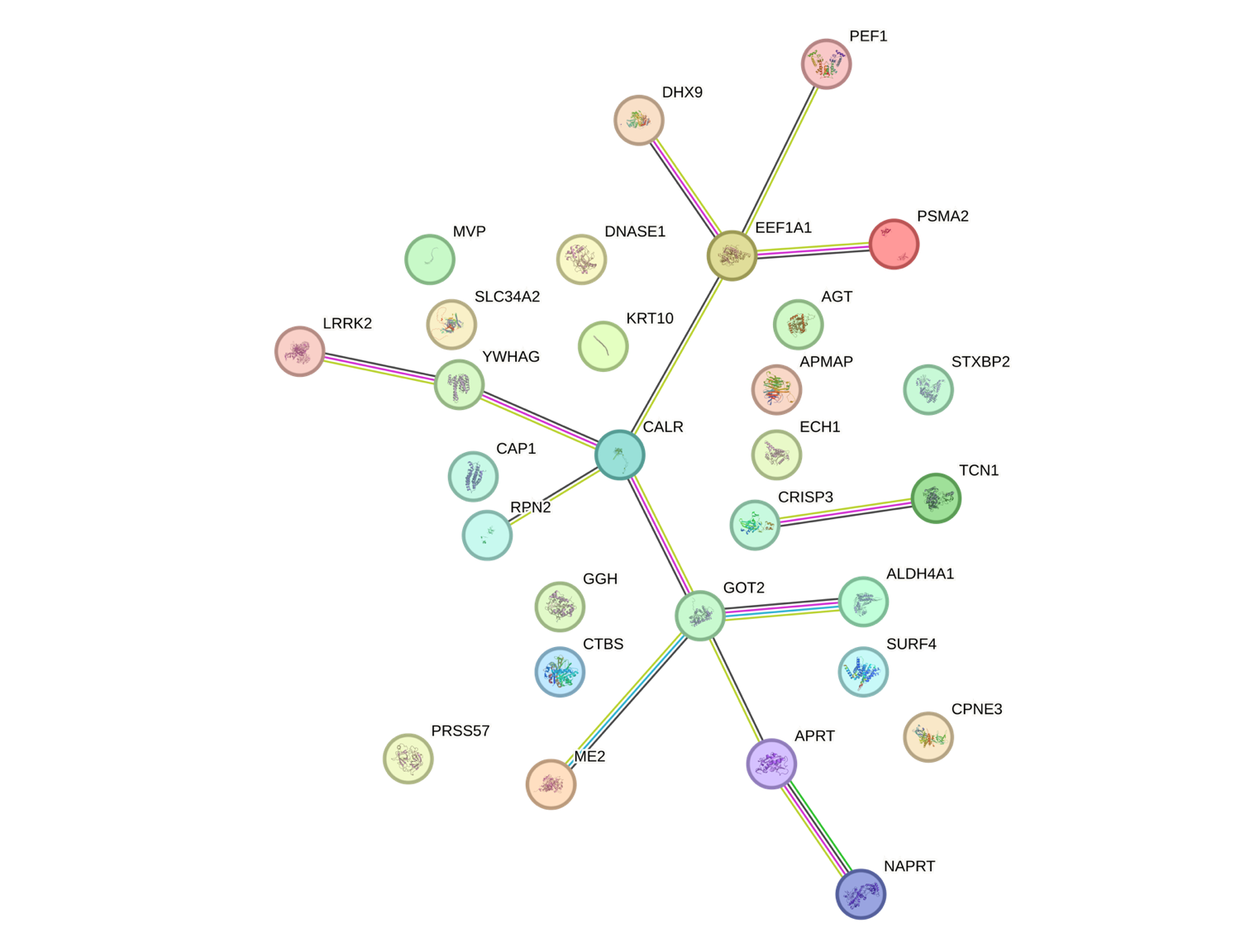

**Supplementary Figure 4: The Protein-Protein interaction Network for the top 30 proteins decreased in abundance when QoL improved.** The STRING database showed that 15/30 (50%) proteins were predicted to have some form of interaction based on known interactions that were either experimentally determined or from curated databases, predicted interactions from gene neighbourhood, gene fusions or gene co-occurance and from other sources such as textmining, co-expression or protein homology.

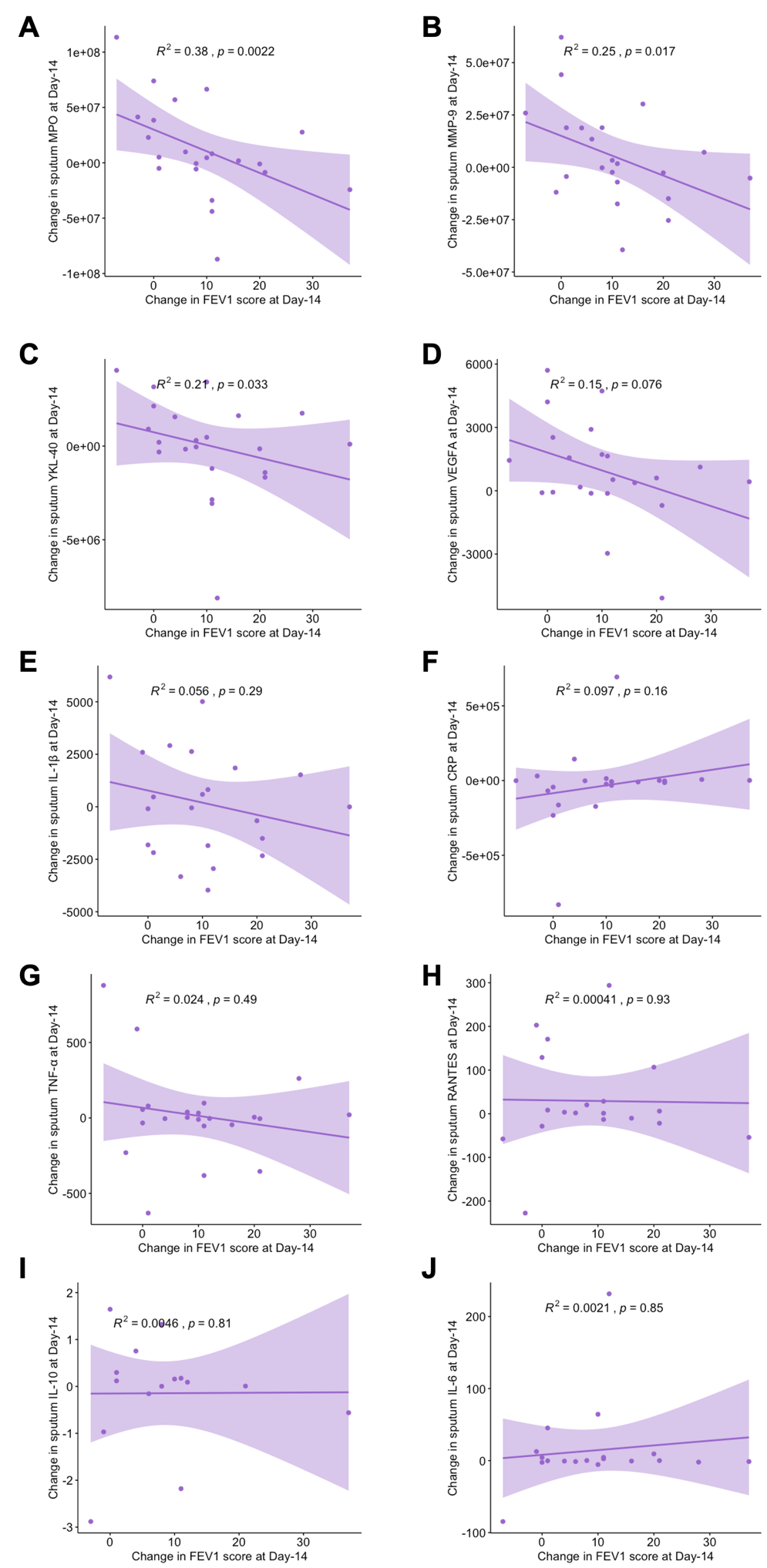

**Supplementary Figure 5: Significant correlations between biomarker reduction and improvement in FEV1 score.** Spearman’s correlation coefficient analysis comparing the changes in host-associated biomarker concentrations between Day-1 and Day-14 to the change in lung function between Day-1 and Day-14 as measured by Forced Expiratory Volume % predicted (FEV1). Specific biomarkers were measured in pg/ml using Mesoscale Discovery assays; MPO (A), MMP-9 (B) and YKL-40 (C) all show significant correlations with an improved FEV1 (% predicted) score. As the FEV1 score improves the concentration of these biomarkers in sputum reduces. The biomarkers VEGF-A (D) and IL-1β (E) also show a reduction in concentration correlated with an improved FEV1 score however this correlation is not significant.

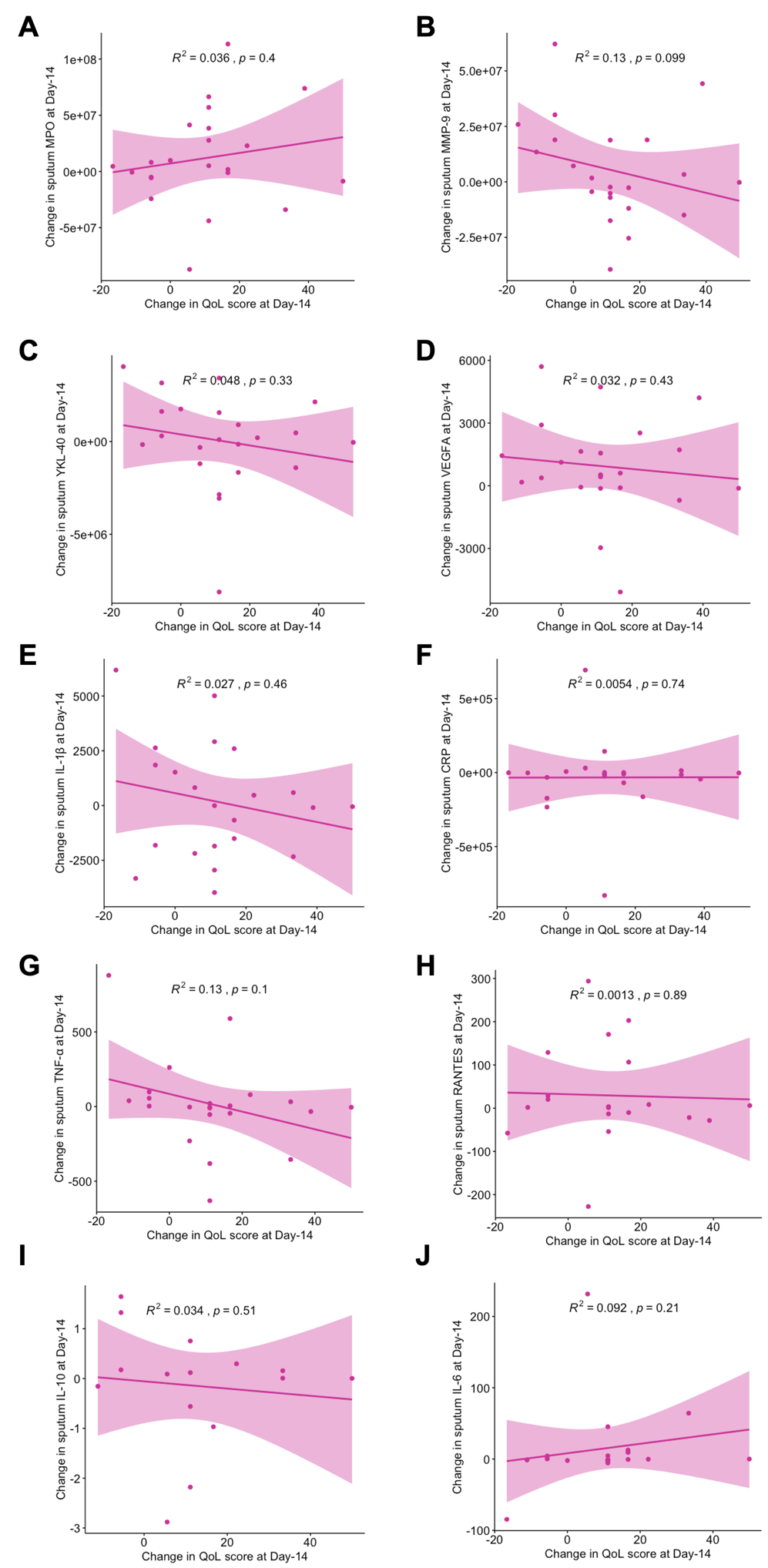

**Supplementary Figure 6: Similar but weaker correlations between reductions in biomarker levels and improvement in CFQ-R score**. Spearman’s Rank Correlation coefficient analysis comparing a change in host-associated biomarker concentration between Day-1 and Day-14 to the change in Quality of Life score between Day-1 and Day-14 as determined by achievement of minimum clinically important difference (MCID) in the CFQ-R Respiratory Domain (QoL). Specific biomarkers were measured in pg/ml using Mesoscale Discovery assays; whilst MMP-9 (B), YKL-40 (C), VEGF-A (D) and IL-1β (E) still show a correlation between reduction in concentration and improved QoL score only MMP-9 (B) is approaching significance with this outcome measure. MPO (A) has a slight positive correlation associated with an increase in concentration when compared to QoL improvement.
